# Targeted knockout of *CYP79A1* reduces cyanogenic potential in grain sorghum

**DOI:** 10.64898/2026.02.23.707491

**Authors:** Evan D. Groover, Jianqiang Shen, Kiflom Aregawi, Sophie Li, Shahar Schwartz, Brian J. Staskawicz, Peggy G. Lemaux, David F. Savage

## Abstract

*Sorghum bicolor* is a climate-resilient C4 crop used for food, forage, and bioenergy, but broader adoption is constrained by accumulation of the cyanogenic glucoside dhurrin, which can release toxic hydrogen cyanide (HCN) upon tissue damage. Dhurrin levels are especially high in juvenile tissues, creating risk for grazing animals and limiting use in mixed crop-livestock systems. Here, we establish a CRISPR-Cas9 genome-editing strategy targeting *CYP79A1*, which catalyzes the first committed step in dhurrin biosynthesis, in the elite grain sorghum inbred RTx430, yielding transgene-free lines with stable, heritable reduction in cyanogenic potential across early vegetative development. Homozygous *cyp79a1* knockouts were negligibly cyanogenic, whereas heterozygous plants exhibited approximately half the cyanogenic potential of unedited nulls. Consistent with established livestock grazing guidelines, only homozygous knockouts fell below thresholds considered hazardous for incidental grazing. This work establishes *CYP79A1* as a practical and heritable genome-editing target for reducing sorghum cyanogenesis and provides a clear path for deployment of low-cyanogenic alleles in elite breeding backgrounds.

## Main Text

*Sorghum bicolor* is a versatile and climate-hardy C4 grass grown globally for food, forage, and bioenergy. In sub-Saharan Africa and South Asia, the expansion of sorghum-based agriculture is closely linked to rural food security and income growth, particularly in drought-prone regions. Despite its importance to smallholder livelihoods, broader adoption of sorghum is constrained by the accumulation of the cyanogenic glucoside dhurrin, a potent anti-herbivory metabolite that is rapidly hydrolyzed to toxic hydrogen cyanide (HCN), causing tissue damage upon ingestion by grazing animals (Gleadow and Møller, 2014). Dhurrin accumulates to particularly high levels in juvenile tissues (Ohio State University Extension, 2021), which are commonly consumed by animals, and its presence in both grain and forage sorghum presents an adoption barrier for many smallholder and subsistence farmers.

Dhurrin biosynthesis is mediated by a metabolon comprising two cytochrome P450 enzymes, CYP79A1 and CYP71E1, which sequentially convert L-tyrosine to *p*-hydroxymandelonitrile, and the UDP-dependent glucosyltransferase UGT85B1, which stabilizes this otherwise labile cyanohydrin through glucosylation to form dhurrin (**Fig. 1A**) (Laursen *et al*., 2016). Studies using chemically mutagenized sorghum lines have established essential roles for each enzyme in this pathway. Loss-of-function mutations in *UGT85B1* cause severe pleiotropic defects consistent with cyanohydrin-mediated autotoxicity (Blomstedt *et al*., 2016). In contrast, missense or nonsense mutations in *CYP79A1* abolish dhurrin production with only modest impacts on early growth and minimal effects on biomass accumulation or agronomic performance in mature plants, consistent with the proposed role of dhurrin as a transient nitrogen reservoir in developing seedlings (Blomstedt *et al*., 2012; Sohail *et al*., 2020). To date, deployment of *CYP79A1* mutant alleles has been limited to a small number of forage and grain sorghum backgrounds. Because elite grain sorghum cultivars can retain substantial cyanogenic potential, reducing cyanogenic potential in grain sorghum addresses a persistent constraint in mixed crop-livestock systems. Here, we establish a genome-editing strategy to reduce or eliminate cyanogenesis in the grain sorghum inbred RTx430 by targeting *CYP79A1*.

**Figure 1.**
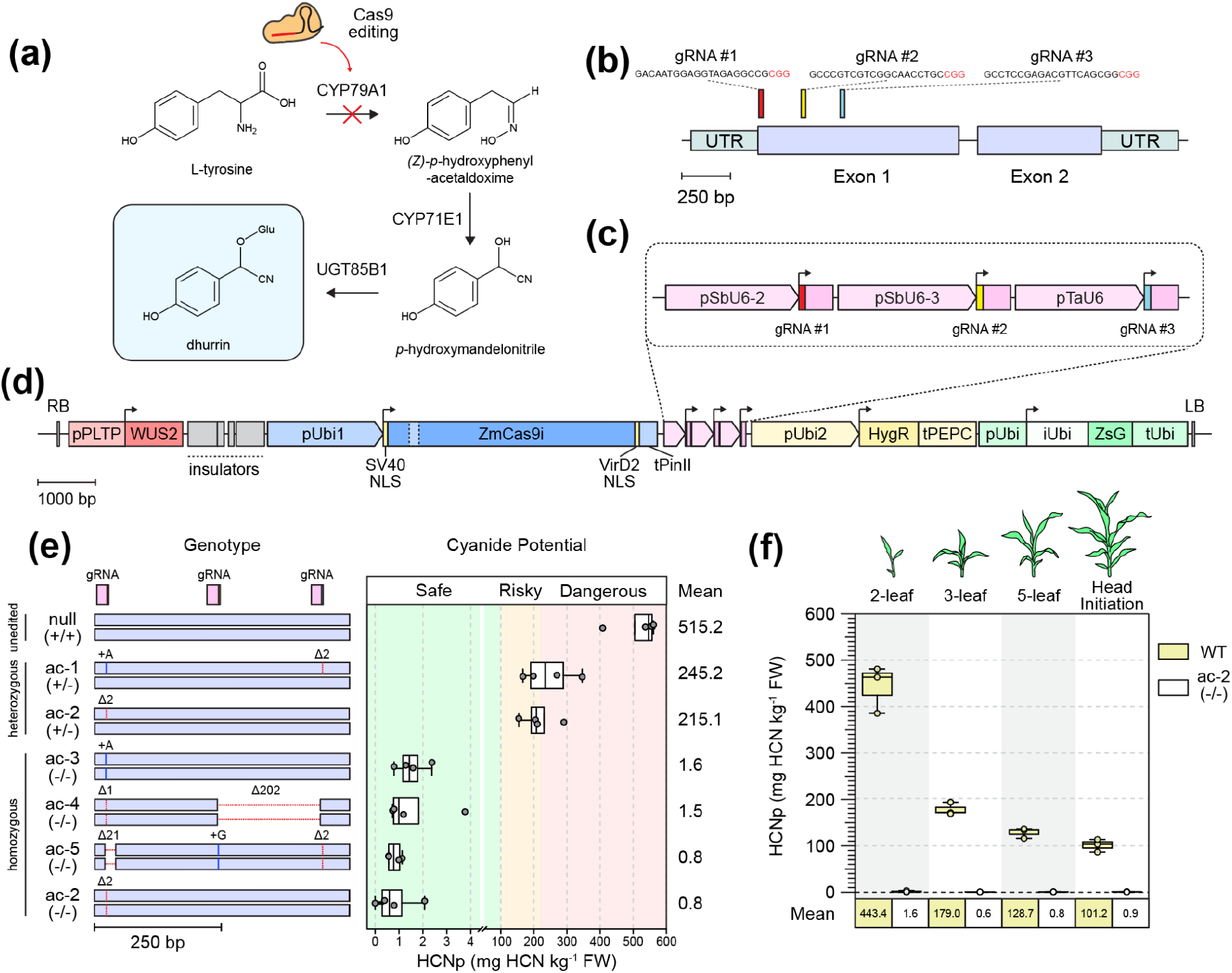
Cas9-mediated editing of *CYP79A1* abolishes sorghum cyanogenicity. **(a)** Enzymatic pathway for dhurrin biosynthesis from L-tyrosine. *CYP79A1* catalyzes the first committed step and was targeted for Cas9-mediated genome editing. **(b)** Gene model of *CYP79A1* showing untranslated regions (UTRs; green) and exons (purple). Three guide RNAs (gRNA #1–3) targeting exon 1 are indicated; protospacer sequences are shown in black with the adjacent PAM motif (NGG) highlighted in red. **(c)** gRNA expression cassettes used for multiplex editing. gRNA #1 is driven by the sorghum *U6-2* promoter, gRNA #2 by the sorghum *U6-3* promoter, and gRNA #3 by the wheat *U6* promoter. **(d)** T-DNA architecture of the binary vector used for transformation. The T-DNA contains a maize codon-optimized Cas9 (ZmCas9i) with two nuclear localization signals (NLS), three guide RNA (gRNA) expression cassettes, a hygromycin resistance marker (hygromycin B phosphotransferase, HygR), a ZsGreen1 fluorescent reporter (ZsG), and the morphogene *WUSCHEL2* (*WUS2*). ZmCas9i is driven by the maize ubiquitin 1 promoter (pUbi1), HygR by the switchgrass ubiquitin 2 promoter (pUbi2), and ZsGreen1 by the sorghum ubiquitin promoter (pUbi). Right and left T-DNA borders (RB and LB) delineate the transferred region. **(e)** Genotypes and cyanogenic phenotype of stable T_1_ lines. Left, haplotype-resolved edited alleles for each line across the targeted region of *CYP79A1*, annotated as heterozygous (+/−) or homozygous (−/−) for edits; an unedited T_1_ control is shown (null, +/+). Right, cyanide potential (HCNp; mg HCN kg^−1^ fresh weight, FW) measured from n = 4 biological replicates per genotype. Background shading indicates livestock grazing risk thresholds (green, safe; orange, risky; red, dangerous) (Ohio State University Extension, 2021). Boxplots show the median (center line) and interquartile range (IQR) (box); whiskers extend to the most extreme values within 1.5× the IQR, and points represent individual biological replicates (outliers beyond the whiskers, if present, are shown as points). The mean HCNp for each genotype is reported at right. **(f)** Developmental dependence of cyanogenic potential in wild-type (WT) and homozygous *cyp79a1* knockout plants from the ac-2 line (e). HCNp was measured at the indicated developmental stages (2-leaf, 3-leaf, 5-leaf, and head initiation) from most recently expanded leaves. Boxplots are defined as in (e) with *n* = 3 biological replicates per genotype; mean values are shown below each group.

Using the pGL222 binary T-DNA system described previously (Shen *et al*., 2025), we targeted the first exon of *CYP79A1* using three evenly spaced guide RNAs (**Fig. 1B**) to increase the likelihood of producing both small insertion-deletions (indels) as well as larger deletions spanning multiple cut sites. Guide RNAs were selected with high predicted on-target activity and minimal off-target potential (**Methods**). Guide RNAs were expressed monocistronically under distinct monocot U6 promoters (**Fig. 1C**) together with an intronized, maize-codon-optimized Cas9. The T-DNA also contained a hygromycin resistance marker for tissue culture selection, the morphogenic gene *WUSCHEL2* to enhance transformation efficiency (Aregawi *et al*., 2022; Shen *et al*., 2025), and the fluorescent marker *ZsGreen1* to facilitate identification of transgenic and transgene-free segregants (**Fig. 1D**).

The final T-DNA construct was transformed into a total of 205 sorghum RTx430 immature embryos (IEs) across two independent rounds of transformation (Aregawi *et al*., 2022). After initial transformation, IEs were recovered in co-cultivation medium for 6 days at 28°C, transferred to resting media for one week at 28°C, moved to embryo maturation media (EMM) for 4-8 weeks with hygromycin selection, and finally transferred to rooting media without selection for root establishment. After two weeks in hygromycin selection in embryo maturation media, explants had an 80.5% embryo transformation efficiency as assessed by fluorescence microscopy (**Supplementary Fig. 1A**).

A subset of 42 regenerated explants were transferred to soil as independent transformants, 40 of which (95.2%) flowered and produced seed (**Supplementary Fig. 1B**). All explants were sequenced to confirm transgene integration and screened for *CYP79A1* editing. Of these, 30 T_0_ transformants showed detectable signals of editing in the seventh fully expanded leaf as assessed by Sanger sequencing using Inference of CRISPR Edits (ICE) analysis, while 12 showed no editing signal (**Supplementary Fig. 1C**). Among reproductively viable edited transformants, efficiencies ranging from 14 to 100% of edited alleles were observed (likely resulting from a heterogeneous mixture of edited and unedited alleles within a leaf sample), with an average editing efficiency of 70.46% (**Supplementary Fig. 1C**). To study the effect of chimeric editing on T_0_ cyanogenesis, a subset of 13 mature T_0_ plants were screened for their cyanogenic potential (HCNp) by a modified picrate assay (Bradbury *et al*., 1999), amended with a dhurrin-hydrolyzing enzyme buffer (Gleadow *et al*., 2012), performed on the seventh fully expanded leaf at maturity. We observed that the CRISPR knockout score was strongly negatively correlated (Pearson’s r = −0.942; R^2^ = 0.887) with cyanide production (**Supplementary Fig. 1D**), indicating that successful editing is quantitatively linked to the depletion of foliar cyanogenic potential. T_0_ transformants with high (>90%) knockout scores were advanced to T_1_, where segregants were screened for absence of ZsGreen1 fluorescence, absence of PCR-amplifiable T-DNA region, and presence of on-target edits. Multiple independent edited, T-DNA-free lines were recovered and advanced for subsequent phenotypic and genotypic analysis.

Genotyped T_1_ plants representing five independent edited backgrounds were screened for cyanogenic potential as homozygotes, heterozygotes, or both, and compared to an unedited null T_1_ control (**Fig. 1E**). Plants were assayed at the two-leaf seedling stage, when dhurrin accumulation is near maximal. Homozygous *cyp79a1* knockouts were negligibly cyanogenic (0.8–1.6 mg HCN kg^−1^ fresh weight), whereas heterozygous plants exhibited approximately half the cyanogenic potential of unedited nulls (**Fig. 1E**). Variation among heterozygous T_1_ individuals likely reflects partial CYP79A1 dosage, but might also be influenced by minor variability in assay performance or tissue sampling. Consistent with established livestock grazing guidelines (Ohio State University Extension, 2021), only homozygous knockouts fell below thresholds considered hazardous for incidental grazing. Depletion of cyanogenic potential was stable across early vegetative development, and a representative homozygous knockout line maintained negligible cyanogenic capacity relative to wild-type controls throughout this period (**Fig. 1F**).

From nine T_0_ transformants exhibiting >90% editing, a total of 14 edited alleles (and two wild-type nulls) were recovered as homozygous T_1_ lines (**Supplementary Fig. 2A**). Based on these outcomes, guide RNA #1 was the most active of the three guides tested, accounting for 88.9% of all edited T_1_ haplotypes and present in all recovered edited alleles (**Supplementary Fig. 2B**). Given its high efficiency and conserved sequence within the early coding region of *CYP79A1* exon 1, this guide represents a strong candidate for deployment in additional sorghum cultivars. As potential off-target edits were not systematically evaluated in this study, further segregation and breeding may be required to separate unintended edits from the desired *CYP79A1* alleles.

Together, these results demonstrate that *CYP79A1* knockout can effectively ablate cyanogenic potential in grain sorghum across a physiologically relevant developmental window, supporting its utility as a practical and heritable genome-editing target for reducing cyanogenic risk in elite grain sorghum cultivars.

## Figures

**Supplementary Figure 1.**
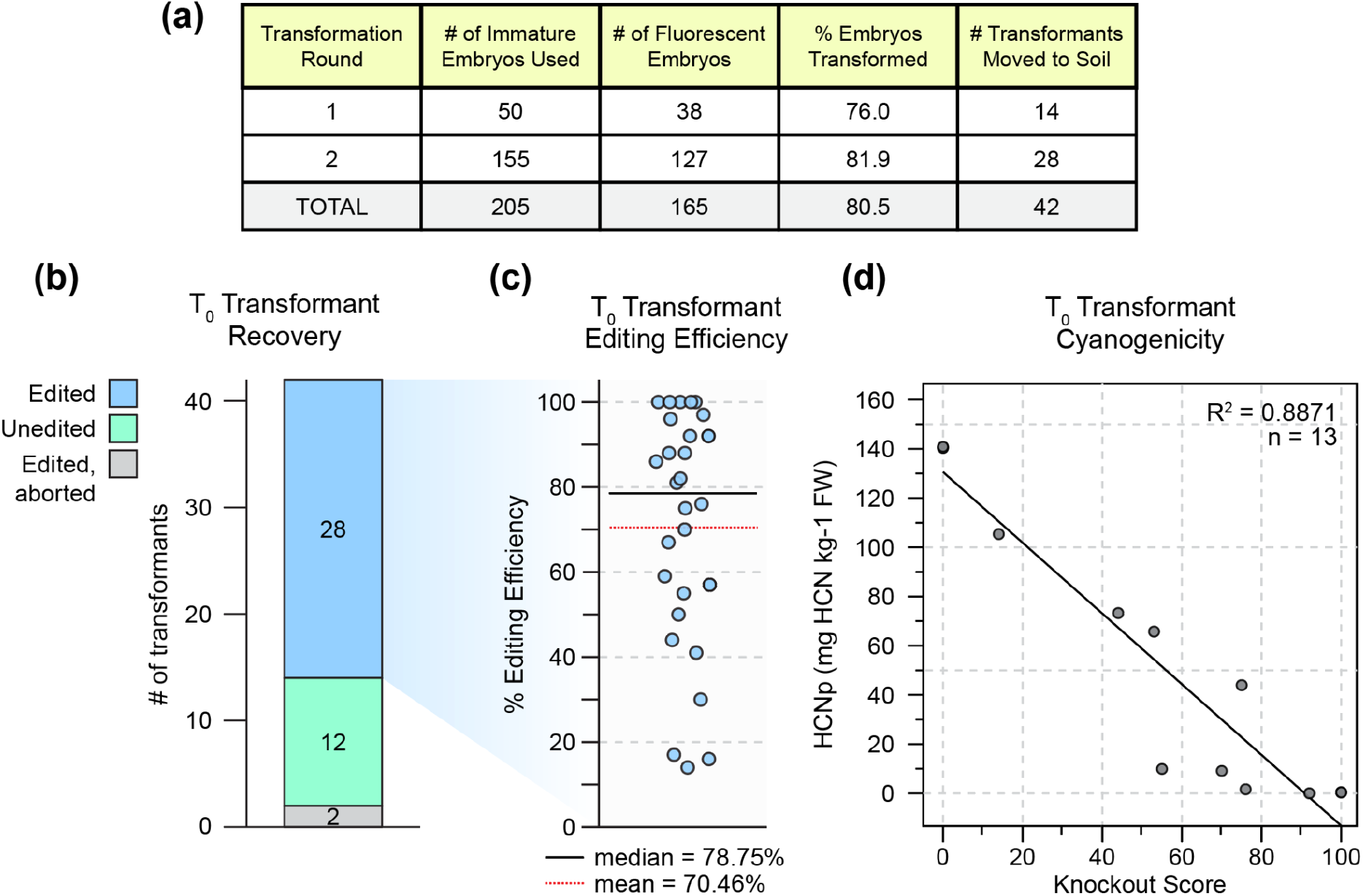
Transformation, genotyping, and screening of T_0_ *CYP79A1* mutants. **(a)** Summary of embryo transformation efficiency across two independent rounds of *Sorghum bicolor* RTx430 immature embryo transformation. Embryo transformation percentage was calculated as the number of independent fluorescent regenerants (not counting subcultures) exhibiting stable fluorescent marker expression at four weeks post-transformation divided by the total number of embryos inoculated. Of 165 fluorescent embryos, 42 independent transformants were transferred to soil and advanced to greenhouse growth. **(b)** Recovery and preliminary genotyping outcomes for T_0_ sorghum transformants assessed for *CYP79A1* editing in the seventh fully expanded leaf. Editing status was determined by Sanger sequencing across the *CYP79A1* target region. “Edited” (blue) indicates detection of one or more indels within the target window; “Unedited” (green) indicates only wild-type sequence was detected; “Edited, aborted” (grey) indicates transformants that did not survive to flowering and could not be phenotyped further. **(c)** Editing efficiencies of all edited T_0_ lines from **(b)**, quantified by Sanger sequencing and ICE analysis. Editing efficiency represents the inferred fraction of edited alleles per line, measured as editing score in ICE. Each point corresponds to a single edited T_0_ event (n = 28). Median (solid black line) and mean (red dashed line) values are indicated. **(d)** Relationship between inferred knockout score derived from ICE analysis and cyanogenic potential in T_0_ plants. Cyanogenic potential (HCNp; mg HCN kg^−1^ FW) was measured from the seventh fully expanded leaf of mature plants and plotted against the knockout score, defined as the inferred fraction of alleles containing frameshift mutations or deletions >21 bp. Each point represents an individual T_0_ plant (*n* = 13). The solid line indicates a least-squares linear regression; coefficient of determination (r^2^) is shown.

**Supplementary Figure 2.**
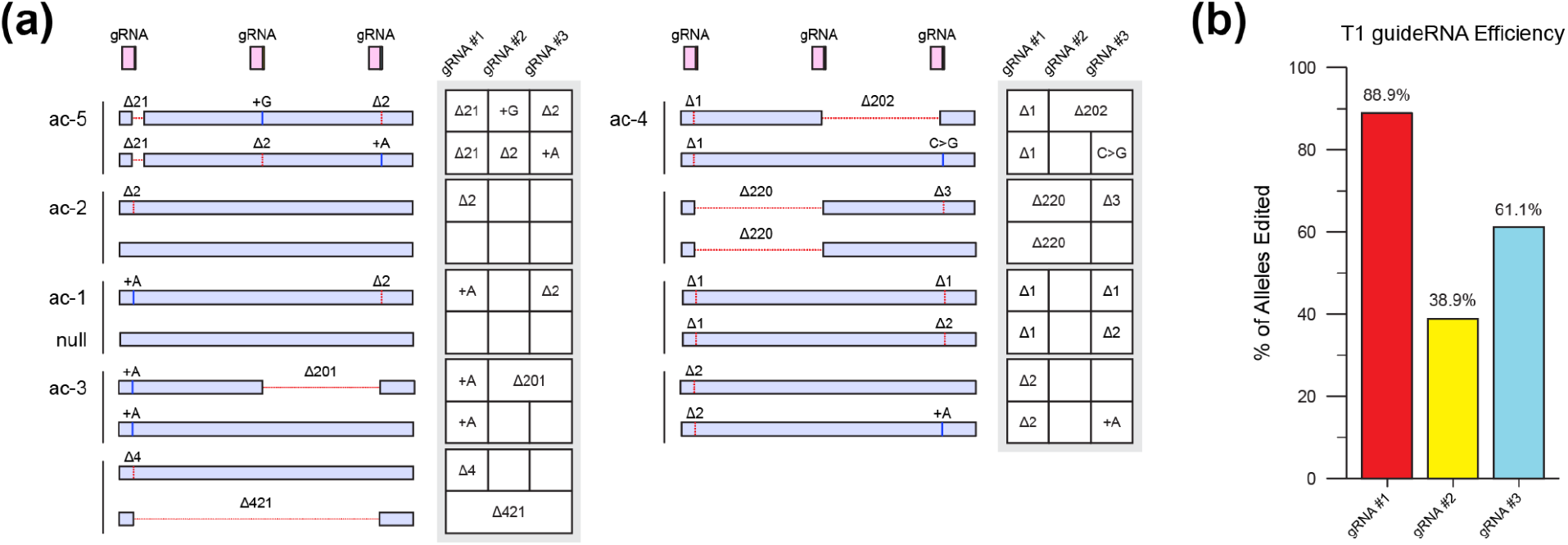
Heritable editing outcomes of *CYP79A1* mutants. **(a)** Haplotype-resolved *CYP79A1* alleles recovered from homozygous edited T_1_ plants. For each T_1_ individual, both alleles are shown schematically, with deletions (Δ), insertions (+), and substitutions annotated relative to the wild-type sequence. Red dashed lines indicate large deletions spanning the multiple guide RNA (gRNA) target sites, which are indicated by pink boxes. Individual alleles were PCR-amplified and sequenced by Sanger sequencing from T_1_ segregants. Vertical bars group lines derived from a common edited T_0_ parent, and line identifiers at left indicate individuals selected for downstream phenotypic analyses (Fig. 1). Summary matrices at right indicate the specific allelic outcomes observed at each gRNA target site for each line. **(b)** Inheritance frequency of edits at each gRNA target site across T1 lines. Bars indicate the percentage of genotyped T1 lines carrying at least one edited allele at the corresponding gRNA target site, calculated across all homozygous edited T1 individuals.

## Methods

### Plant material and growth conditions

Seeds of *Sorghum bicolor* genotype RTx430 were obtained from the Germplasm Resources Information Network (GRIN, USDA-ARS; PI 655998). T_0_ transformant plants were cultivated in 3-gallon pots containing SuperSoil (Rod McLellan Company) in the UC Berkeley Oxford Research Unit South Greenhouse under controlled conditions of 30°C/24°C (day/night) with a 16-hour light/8-hour dark photoperiod. T_1_ plants were cultivated in 3.5-inch pots in a growth chamber at 28°C, 100 μmol photons m^−2^ s^−1^ light intensity, a 16-hour light/8-hour dark photoperiod, and 80% relative humidity.

### gRNA and CRISPR Construct Creation

Candidate target sequences within *CYP79A1* were identified in the *S. bicolor* cv. RTx430 v2.1 genome (Phytozome; https://phytozome-next.jgi.doe.gov/info/SbicolorRTx430_v2_1). Three guide RNAs targeting the first exon of *CYP79A1* (SbiRTX430.01G012200) were designed using the Benchling CRISPR tool. Guides were selected based on high predicted on-target activity and low predicted off-target potential. Guide RNAs were spaced approximately 200 bp apart within the coding sequence to increase the likelihood of large (hundreds of bases), loss-of-function deletions between cut sites. Targeted regions were PCR-amplified from genomic DNA and verified by Sanger sequencing prior to cloning. Notably, guide RNAs #1 and #3 have no predicted complete genomic off-target sites in the sorghum pangenome, but guide RNA #2 has one homologous off-target in some sorghum cultivars in another cytochrome P450 gene (SbiRTX430.10G183100), which is predicted to act orthogonally in phenylacetaldoxime biosynthesis. Although prospective off-target sequences were not sequenced in this study, targeted or whole-genome sequencing to detect genomic off-target mutations should be performed for regulatory assessment and field deployment. Notably, *cyp79a1* mutant lines displayed growth rates and overall growth habits indistinguishable from unedited control plants.

Guide RNAs were cloned into the pGL222 binary vector using a modified shuttle vector system described previously (Shen *et al*., 2025). Guide RNA spacers were introduced into a modified pGL198 shuttle vector (Addgene #180472) containing a tandem array of monocot U6 promoters (Sorghum *U6*.*2*.*3*, Sorghum *U6*.*3*.*1*, and *Triticum aestivum* U6), each monocistronically driving a guide RNA scaffold. The shuttle vector also contained an intronized maize codon-optimized Cas9 driven by the maize ubiquitin-1 promoter and terminated by the potato PINII terminator. Spacer sequences were introduced by Gibson Assembly (NEBuilder HiFi DNA Assembly Master Mix, NEB E2621L). Shuttle vectors were transformed into chemically competent TOP10 *E. coli* and fully sequenced to confirm assembly. Verified shuttle vectors were recombined into pGL222 using Gateway LR Clonase II (Invitrogen 11791020), and final constructs were sequence-verified.

### *Agrobacterium*-mediated transformation

Binary constructs were introduced into *Agrobacterium tumefaciens* strain LBA4404 Thy^−^ by electroporation. Transformants were selected on YEP medium supplemented with 100 mg L^−1^ thymidine, 50 mg L^−1^ gentamicin, and 50 mg L^−1^ spectinomycin. Immature embryos were harvested 12–14 days post-anthesis and co-cultivated with *Agrobacterium* suspension (OD_550_ = 0.7) prepared in PHI-I medium supplemented with 0.005% (v/v) Silwet L-77 and 0.2 mM acetosyringone. Embryos were co-cultivated at 28°C for one week, transferred to resting medium for one week at 28°C in the dark, and then placed on embryo maturation medium containing 15 mg L^−1^ hygromycin for 4–6 weeks until regeneration. Regenerants were transferred to rooting medium and subsequently to soil. All media conditions are as described in (Aregawi *et al*., 2022; Shen *et al*., 2025).

### Visualization of T-DNA integration

T-DNA integration and segregation were monitored using the ZsGreen1 fluorescent reporter. Fluorescence was assessed weekly using a Zeiss SteREO Lumar.V12 epifluorescence stereomicroscope equipped with a QImaging Retiga SRV camera. Embryo transformation efficiency was calculated as the number of independent fluorescent regenerants (not counting subcultures, i.e. where more than one transformant came from a single embryo) divided by the total number of embryos inoculated.

### DNA Extraction

Genomic DNA was extracted using a Chelex-based protocol. Leaf tissue (~1 cm^2^) was homogenized in 280 μL of 10% (w/v) Chelex® 200–400 mesh resin (Bio-Rad 95621) prepared in 5 µM Tris-HCl (pH 8.0). Samples were vortexed, incubated at 95°C for 5 min, vortexed again, and centrifuged at maximum speed for 1 min. Supernatant was used directly for PCR amplification.

### Genotyping transformants and quantifying editing

T-DNA presence was assessed using primers spanning a 1.5-kb region within the T-DNA region and starting immediately downstream of the T-DNA left border. *CYP79A1* edits were genotyped using primers amplifying a ~1 kb region encompassing all three guide RNA target sites. Amplicons were Sanger sequenced, and editing outcomes were quantified using EditCo ICE analysis (ice.editco.bio). ‘Editing efficiency’ refers to the percentage of alleles containing any guide-proximal edit, whereas ‘knockout score’ refers to alleles containing frameshift mutations or deletions >21 bp. T_1_ seedlings were screened for fluorescence using a Leica M205C stereomicroscope, and non-fluorescent plants were genotyped using leaf punches collected at the time of cyanogenic analysis.

### Cyanide quantification

Cyanogenic potential was quantified using a modified sodium picrate assay (Bradbury *et al*., 1999) optimized for complete enzymatic hydrolysis of dhurrin. Foliar cyanogenic glucoside detection kits were obtained from the Australian National University (Kit E; https://biology.anu.edu.au/research/divisions/konzo-prevention-unit/kits-posters) and adapted for quantitative analysis using a β-glucosidase degradation buffer commonly employed for dhurrin hydrolysis (Gleadow *et al*., 2012).

Mid-leaf tissue was harvested from the second, third, fifth, or seventh fully expanded leaf (i.e., no further blade elongation) prior to the emergence of the subsequent leaf. For all comparisons, plants were sampled based on matched developmental stage (leaf number and expansion status), such that knockout and control plants were assayed at equivalent physiological stages, even when this required sampling at slightly different chronological ages. Leaf tissue was sliced into ~2 mm strips using a safety razor and thoroughly ground with a mortar and pestle until complete homogenization was achieved. All tissue processing steps were optimized to ensure maximal cyanide release within the reaction vessel, and mechanical disruption was performed as rapidly as possible to minimize premature volatilization or loss of hydrolyzed cyanide prior to measurement.

For each sample, 100 mg of homogenized fresh tissue was weighed and transferred to a sealed reaction vial containing 1 mL of β-glucosidase degradation buffer composed of β-glucosidase (2 mg mL^−1^; Sigma G0395) dissolved in 0.1 M trisodium citrate buffer (pH ~5.5). A sodium picrate indicator strip was suspended within each vial. Negative control vials lacking tissue and positive control vials supplied with the kit were included in all assays. β-glucosidase degradation buffer was prepared fresh for each experiment. Sealed vials were incubated for 24 h at room temperature in the dark to allow complete enzymatic hydrolysis of dhurrin and absorption of released hydrogen cyanide by the indicator strip.

Following incubation, indicator strips were removed and eluted in 5 mL of ddH_2_O in 15-mL culture tubes. Tubes were incubated for 30 min, including 10 min of agitation at 40 rpm on a rotary shaker. Eluates were mixed by pipetting, and absorbance was measured at 510 nm using an Ultrospec 3000 UV/Visible spectrophotometer (Pharmacia Biotech). Absorbance values were normalized to negative controls and multiplied by a factor of 396 to calculate cyanogenic potential, expressed as mg HCN kg^−1^ fresh weight.

Assay accuracy was verified in all experiments using kit-provided positive controls and potassium cyanide (KCN) standards spanning a range of concentrations. For T_1_ phenotyping experiments, leaf tissue from three to four independent plants per genotype was analyzed. For T_0_ phenotyping, samples were obtained non-destructively by collecting leaf hole punches from the seventh fully expanded leaf following transfer to the greenhouse. For all phenotyping, nontransformed (T_0_ and T_1_) or unedited (T_1_) negative control plants were measured in parallel.

## Author Contributions

Conceptualization: EDG, DFS.

Transformation: EDG, SL, JS, KA.

Analysis: EDG, SL, SS.

Writing: EDG.

Visualization: EDG.

Supervision: BJS, PGL, DFS.

Project administration and funding acquisition: DFS.

## Acknowledgements

The authors would like to thank Alicia Quinn, Nicholas Karavolias, Emma Choi, and Jessica Lyons for their advice and assistance in establishing the cyanide measurement assay.

## Data Availability Statement

All data supporting the findings of this study are included in the manuscript and Supplementary Information. Sanger sequencing data, plasmid sequences, and associated analysis files are available from the corresponding author upon reasonable request.

## Ethics / Biosafety Statement

All experimental procedures involving genetically modified plants were conducted in compliance with institutional biosafety regulations and approved containment practices at the University of California, Berkeley. No human or animal subjects were involved in this study.

## References

Aregawi, K., Shen, J., Pierroz, G., Sharma, M.K., Dahlberg, J., Owiti, J., and Lemaux, P.G. (2022) Morphogene-assisted transformation of Sorghum bicolor allows more efficient genome editing. Plant Biotechnol. J., 20, 748–760.

Blomstedt, C.K., Gleadow, R.M., O’Donnell, N., Naur, P., Jensen, K., Laursen, T., et al. (2012) A combined biochemical screen and TILLING approach identifies mutations in Sorghum bicolor L. Moench resulting in acyanogenic forage production: Acyanogenic forage sorghum plants. Plant Biotechnol. J., 10, 54–66.

Blomstedt, C.K., O’Donnell, N.H., Bjarnholt, N., Neale, A.D., Hamill, J.D., Møller, B.L., and Gleadow, R.M. (2016) Metabolic consequences of knocking out UGT85B1, the gene encoding the glucosyltransferase required for synthesis of dhurrin in Sorghum bicolor (L. Moench). Plant Cell Physiol., 57, 373–386.

Bradbury, M.G., Egan, S.V., and Bradbury, J.H. (1999) Picrate paper kits for determination of total cyanogens in cassava roots and all forms of cyanogens in cassava products. J. Sci. Food Agric., 79, 593–601.

Gleadow, R.M., Møldrup, M.E., O’Donnell, N.H., and Stuart, P.N. (2012) Drying and processing protocols affect the quantification of cyanogenic glucosides in forage sorghum. J. Sci. Food Agric., 92, 2234–2238.

Gleadow, R.M. and Møller, B.L. (2014) Cyanogenic glycosides: synthesis, physiology, and phenotypic plasticity. Annu. Rev. Plant Biol., 65, 155–185.

Laursen, T., Borch, J., Knudsen, C., Bavishi, K., Torta, F., Martens, H.J., et al. (2016) Characterization of a dynamic metabolon producing the defense compound dhurrin in sorghum. Science, 354, 890–893.

Ohio State University Extension (2021) FAQs about cyanide or “prussic acid” poisoning in ruminants.

Shen, J., Aregawi, K., Anwar, S., Miller, T., Groover, E., Rajkumar, M., et al. (2025) High-frequency sorghum transformation toolkit enhances Cas9 efficiency and expands promoter-editing capability with SpRY. bioRxiv, 2025.01.21.634149.

Sohail, M.N., Blomstedt, C.K., and Gleadow, R.M. (2020) Allocation of resources to cyanogenic glucosides does not incur a growth sacrifice in sorghum bicolor (L.) Moench. Plants, 9, 1791.

